# Characterization of fluorescent proteins with intramolecular photostabilization

**DOI:** 10.1101/2020.03.07.980722

**Authors:** Sarah S. Henrikus, Konstantinos Tassis, Lei Zhang, Jasper H. M. van der Velde, Christian Gebhardt, Andreas Herrmann, Gregor Jung, Thorben Cordes

## Abstract

Genetically encodable fluorescent proteins have revolutionized biological imaging *in vivo* and *in vitro*. Since there are no other natural fluorescent tags with comparable features, the impact of fluorescent proteins for biological research cannot be overemphasized. Despite their importance, their photophysical properties, i.e., brightness, count-rate and photostability, are relatively poor compared to synthetic organic fluorophores or quantum dots. Intramolecular photostabilizers were recently rediscovered as an effective approach to improve photophysical properties. The approach uses direct conjugation of photostablizing compounds such as triplet-state quenchers or redox-active substances to an organic fluorophore, thereby creating high local concentrations of photostabilizer. Here, we introduce an experimental strategy to screen for the effects of covalently-linked photostabilizers on fluorescent proteins. We recombinantly produced a double cysteine mutant (A206C/L221C) of α-GFP for attachment of photostabilizer-maleimides on the ß-barrel in close proximity to the chromophore. Whereas labelling with photostabilizers such as Trolox, Nitrophenyl, and Cyclooctatetraene, which are often used for organic fluorophores, had no effect on α-GFP-photostability, a substantial increase of photostability was found upon conjugation of α-GFP to an azobenzene derivative. Although the mechanism of the photostabilizing effects remains to be elucidated, we speculate that the higher triplet-energy of azobenzene might be crucial for triplet-quenching of fluorophores in the near-UV and blue spectral range. Our study paves the way towards the development and design of a second generation of fluorescent proteins with photostabilizers placed directly in the protein barrel by methods such as unnatural amino acid incorporation.

## 1. Introduction

Fluorescent proteins (FPs) have revolutionized fluorescence imaging of biological systems *in vivo* and *in vitro*. Because they are genetically encoded, they allow the tethering of a natural light-emitting protein chromophore to any protein of interest^1–3^. Since there are no other fluorescent tags with these properties, the impact of FPs for biological research cannot be overemphasized^1, 3–5^. Despite their importance, the photophysical properties of FPs, i.e., brightness, count-rate and photostability^6–8^, are relatively poor compared to synthetic organic fluorophores^9^ or quantum dots^10–11^. Extensive research has been done over the past decades to improve the photophysical properties of FPs^12^. These studies have resulted in numerous FP-variants^13–15^ with useful chemical and photophysical properties, such as variants optimized for fast folding^16–17^, photoswitching^18^, and brigthness^8, 19–20^, or for functions such as pH sensing^21^. Yet, there are no FPs with photophysical properties that can compete with synthetic dyes in terms of brightness and photostability^6^.

Intramolecular triplet-state quenchers were recently rediscovered as an attractive approach for photostabilization in various fluorescence applications^22–23^. The approach developed in the 1980s^24–25^ uses direct conjugation of photostablizing compounds such as triplet-state quenchers or redox-active substances to a fluorescent reporter (typically a synthetic organic fluorophore), thereby creating high local concentrations of photostabilizer around the fluorophore^27^. As illustrated in Figure 1, this improves the photophysical properties of organic dyes such as Cy5 in bulk and single-molecule investigations *via* intramolecular quenching of triplet or radical states, or; photo-induced electron transfer reactions (mediated in the concrete example by the nitrophenylalanine (NPA) group; data from ref ^27^).

**Figure 1.**
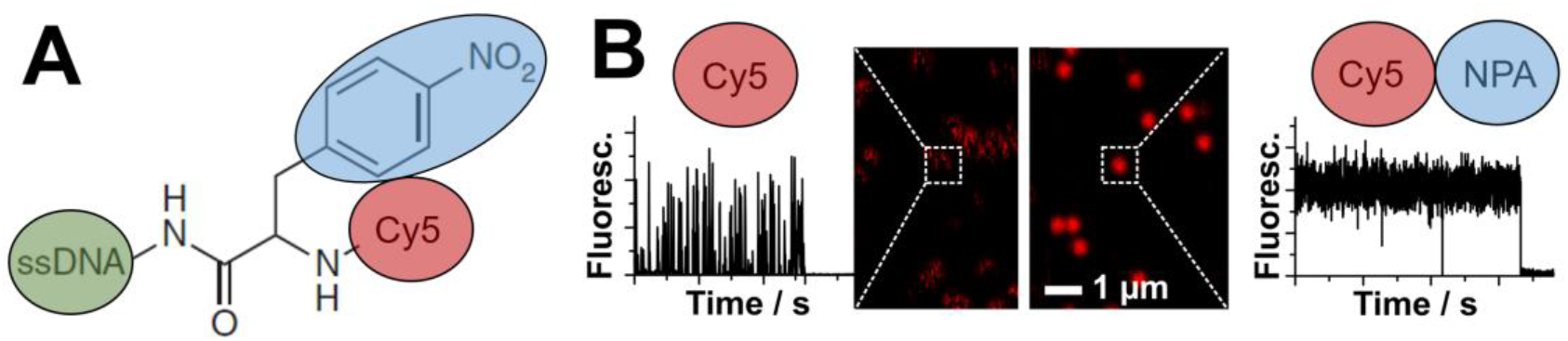
A) Structure of a self-healing organic NPA-Cy5 fluorophore on an oligonucleotide structure (ssDNA). B) Experimental demonstration of photostability increases of Cy5 that are simultanously coupled to a biomolecule (left) and to a photostabilizer (right). Analysis of single-molecule fluorescence microscopy data shows temporal behaviour of fluorescence emission of ‘self-healing’ fluorophore and confocal scanning images and time traces from self-healing Cy5 fluorophores on oligonucleotides. Data reprinted from ^27^.

Such a strategy obviates the need for complex buffer systems, and makes these dyes with intramolecular photostabilization “self-healing”, and thus compatible with diverse biological systems^22–23, 26–29^. This is a particular advantage in situations in which the fluorescent dye is inaccessible to exogenously added stabilizers (e.g., when contained in certain biological cell-compartments^30^). Based on new mechanistic insights^31–32^, there has been exciting progress on the optimization of the photostabilization efficiencies in self-healing dyes^30, 33–35^, the development of bioconjugation strategies for different fluorophore types^27^, photostabilizers and biomolecules^27, 36^, and their new applications in super-resolution^22, 27, 37^, live-cell and single-molecule imaging. All this activity, however, has so far been focused on the major classes of synthetic organic fluorophores including rhodamines^23, 27, 33, 37^, cyanines^22, 27–28, 30, 34–35^, carbopyronines^37^, bophy-dyes^38^, oxazines^36^ and fluoresceins^36^. The recent direct and unambiguous demonstration of the formation of a long-lived chromophore triple state in green fluorescent proteins^39^ suggests that intramolecular photostabilization may be a strategy applicable to fluorescent proteins as well.

The green fluorescent protein (GFP) was discovered by Shimomura et al. in the jellyfish Aequorea victoria (avGFP) in 1962^5^. The 27 kDa protein shows a secondary structure made up of eleven β-strands, two short α-helices and the chromophore in the center. The β-strands form an almost perfect barrel, which is capped at both ends by α-helices^40^. Therefore the para-hydroxybenzylidene-imidazolinone chromophore in the center of the β-barrel is completely separated from exterior^41^. The dimension of the cylinder are 4.2 by 2.4 nm. Proper folding is required for autocatalytic maturation of the chromophore from the amino acids Ser65, Tyr66 and Gly67^41^. GFP shows green fluorescence after excitation in the near UV and blue spectral region. A major and minor absorption peak at 395 nm and 475 nm, respectively, describes the spectral characteristics of GFP. Fluorescence emission occurs either at 503 nm (excitation at 475 nm) or 508 nm (excitation at 395 nm). The two emission peaks belong to two chemically distinct species of the chromophore, namely the anionic form or the neutral phenolate. Excellent summaries of GFP photophysics are provided in refs. ^15, 42–43^.

Here, we introduce an experimental strategy to screen for the effects of covalently-linked photostabilizers on fluorescent proteins. For this, we recombinantly produced a double cysteine mutant (A206C/L221C, Figure S1) of alpha-GFP (F99S/M153T/V163A)^44^ for attachment of photostabilizer-maleimide conjugates. The cysteines did not influence the fluorescence parameters, i.e., spectrum and quantum yield, of the protein and also labelling with cylcooctatetraene (COT), trolox (TX) and a nitrophenyl-group showed negligible effects. Strikingly, we found a substantial increase of photostability upon conjugation to the azobenzene (AB) derivative, 4-phenylazomaleinanil (4-PAM, Figure S1C). Although the mechanism underlying FP-photostabilization by azobenzene remains to be elucidated, our study paves the way towards the development and design of a second generation of fluorescent proteins with photostabilizers placed directly in the protein barrel by methods such as unnatural amino acid incorporation.

## 2. Results

A key obstacle in designing our research was the complex photophysical behavior of FPs, which meant that not only the properties of the chromophore itself, but also factors such as the ß-barrel structure/biochemical state and the specific environment of the proteins had to be considered^45–48^. Although unnatural amino-acid incorporation does present an attractive strategy for the introduction of a photostabilizer into an FP, this route seemed challenging due to low protein expression levels or incorrect protein folding. Therefore, we decided for a strategy where photostabilizers can be covalently linked to GFP via thiol-malemide chemistry (Figure 2A).

**Figure 2.**
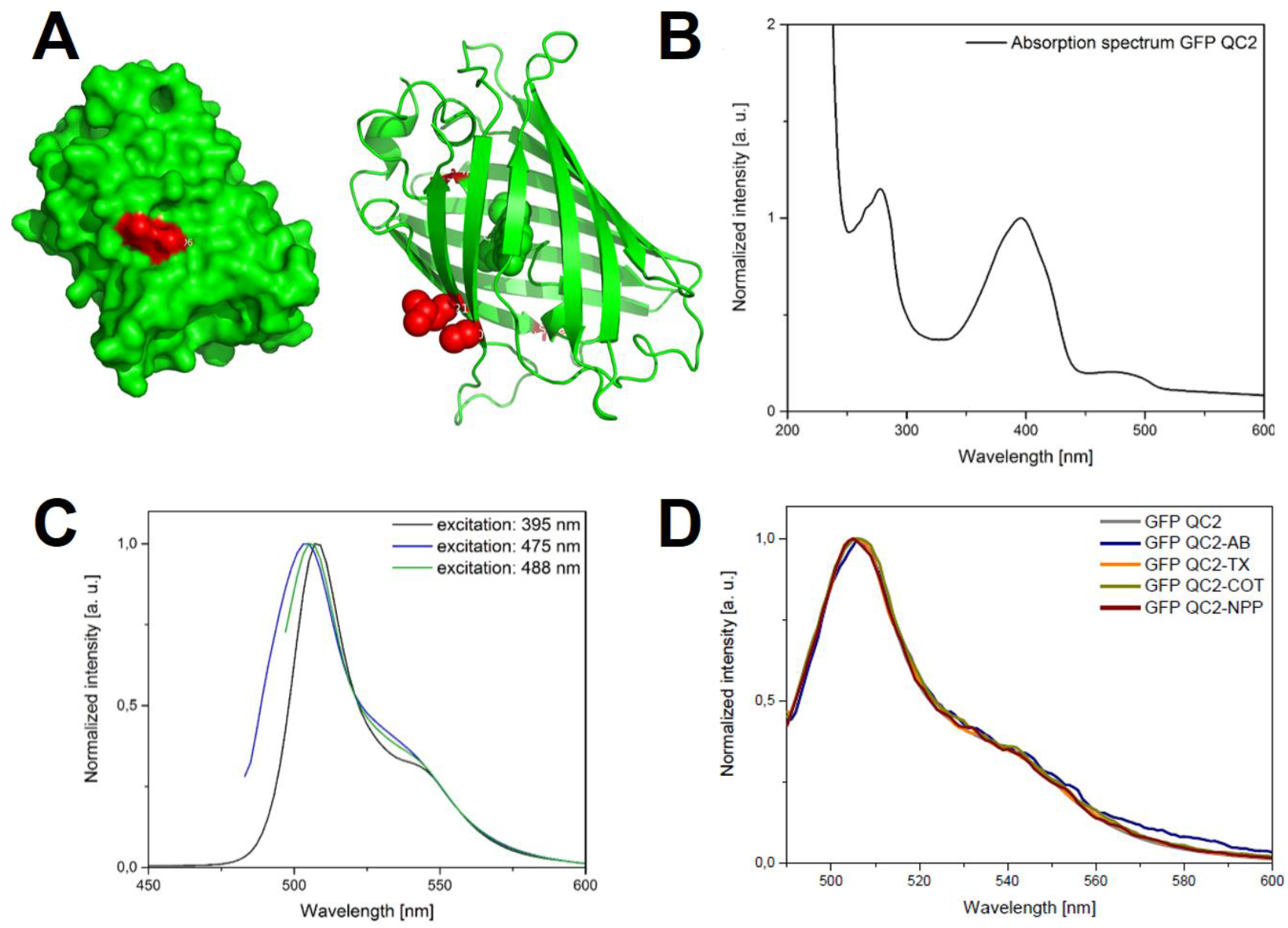
**(A)** Crystal structure of GFP-QC2 indicating residues A206 and L221 in red. These residues were substituted with cysteines in this study for attachment of maleimide photostabilizers. **(B)** Absorbance and **(C)** emission spectra, and **(D)** normalized emission spectra of unlabeled and labeled GFP-QC2.

We produced a double cysteine mutant of α-GFP, a GFP variant with mutations F99S/M153T/V163A as compared to wildtype GFP. We call this variant GFP-QC2 since it additionally contains two solvent-accessible cysteine residues (A206C, L221C, Figure 2A). The side chains of A206 and L221 are directed to the outside of the ß-barrel, and therefore, following cysteine substitution of these residues, and labelling, photostabilizers can be placed outside of the barrel.

The idea was that A206C and L221C (Figure 2A) would be points of attachment for photostabilizers that can affect the chromophore via changes of the protein-barrel^49^ or alternatively via triplet energy-transfer processes using long-lived triplet-states^39^. While the latter are believed to occur more likely via Dexter-processes^22–23^, which would require collisions between FP chromophore and photostabilizer, there is support that certain triplet quenchers might utilize a Förster mechanism^50^. We thus reasoned that intramolecular triplet-quenching in FPs might not strictly require direct contacts between chromophore and stabilizer but proximity. This idea is strongly supported by the observation that FPs can also be influenced by solution-based photostabilizers (Figure S2 and refs. ^51–53^). Tinnefeld and co-workers also demonstrated that EYFP shows a 6-fold enhanced photostability when using dSTORM/ROXS-buffer, i.e., a reducing-oxidizing buffer cocktail, oxygen removal and thiol addition^54^.

α-GFP contains two natural cysteines (C48, C70) which may have potentially interfered with our desired labeling of the barrel using maleimide chemistry. C48 is solvent-accessible, but too far away from the chromophore itself to be useful for photostabilizer attachment and was therefore removed by substitution for a serine residue (Figure S1A). In contrast, C70 is not solvent-accessible in the folded form of GFP, and was therefore not expected to interfere with labeling (Figure S1B). The final construct GFP-QC2 was verified by sequencing to carry the following mutations: C48S/F99S/M153T/V163A/A206C/L221C (Material and Methods & Figure S4).

The absorption and emission properties of GFP-QC2 were analyzed by steady-state spectroscopy methods^27^, and the results of these analysis are given in Figure 2/S3. The spectral characteristics of GFP-QC2 resembled those of α-GFP^55^. The absorption spectrum of GFP-QC2 shows a main peak at ~395 nm (neutral chromophore) and a smaller peak at ~475 nm (anionic chromophore). In the UV range, absorbance by the aromatic amino acids tryptophan, tyrosine and phenylalanine, dominatedand dominate the absorption spectrum giving rise to an additional peak at ~280 nm. An important characteristic of the absorption spectrum was that the ratio of extinction coefficients of GFP-QC2 was slightly below ~1 at 280/395 nm.

Importantly, GFP-QC2 shows a fluorescence spectrum and quantum yield^55^ of 0.81±0.02 (Figure S3) which resemble those of α-GFP. Also the presence or absence of TCEP does not influence the spectra and quantum yield (0.81±0.01), suggesting that cysteine oxidation or di-sulfide bridge formation does not occur in GFP-QC2. We also determined the quantum yield of eGFP to validate our method and found values of 0.63±0.02 and 0.63±0.02 in the absence and presence of TCEP, respectively (Figure S3). All this supports the idea that the cysteines A206C/L221C will provide anchor points for covalent attachment of photostabilizers, but do not influence the photophysics of the FP-chromophore, e.g., by modification of the barrel-structure.

To test for intramolecular photostabilization, we compared the photophysical properties of unlabeled GFP-QC2 with labelled variants carrying the photostabilizers 4-PAM, Trolox (TX), cyclooctatetraene (COT) and nitrophenyl (NPP); see SI for details of photostabilizer synthesis. TX, COT and NPP are photostabilizers that have been extensively used in self-healing dyes due to their triplet-state energy matching with organic fluorophores for Dexter-transfer (COT) or photo-induced electron-transfer (TX, NPP).^22–23, 26–29^ Azobenzene and stilbene, used in the original articles by Lüttke and co-workers for POPOP-dyes are both known as potent quenchers of triplet-states^56^. Since solution-quenching of triplet-states with rate constants up to ~10^10^ M^−1^s^−1^ were observed using azobenzene^56^, this molecule is generally an interesting candidate for both intra- and intermolecular photostabilization. Reasons for not selecting azobenzene earlier on in the development of self-healing dyes may have been caused by its additional ability to induce phototriggered conformational changes (in biological structural such as proteins^57–59^), which require additional control experiments of biochemical function.

Labelling of GFP-QC2 was achieved using a protocol adapted from single-molecule Förster resonance energy transfer experiments^60^ (details see SI: 2. Material and Methods). The labelling of GFP-azobenzene (GFP-AB) was monitored by size exclusion chromatography (Figure 3) via absorbance measurements at 280 nm (Trp/Tyr absorbance of GFP), 320 nm (4-PAM) and 395 nm (GFP chromophore). For GFP-QC2, the 280/395 ratio was just below 1 (Fig. 3A), whereas it was just above 1 for GFP-AB (Fig. 3B). These findings are consistent with the absorption spectrum of GFP-QC2 in Figure 2. A clear indication for labelling of GFP with the azobenzene-derivative 4-PAM is an absorbance increase at 320 nm (Fig. 3A vs. 3B; see 4-PAM absorbance spectrum in Figure S1).

**Figure 3.**
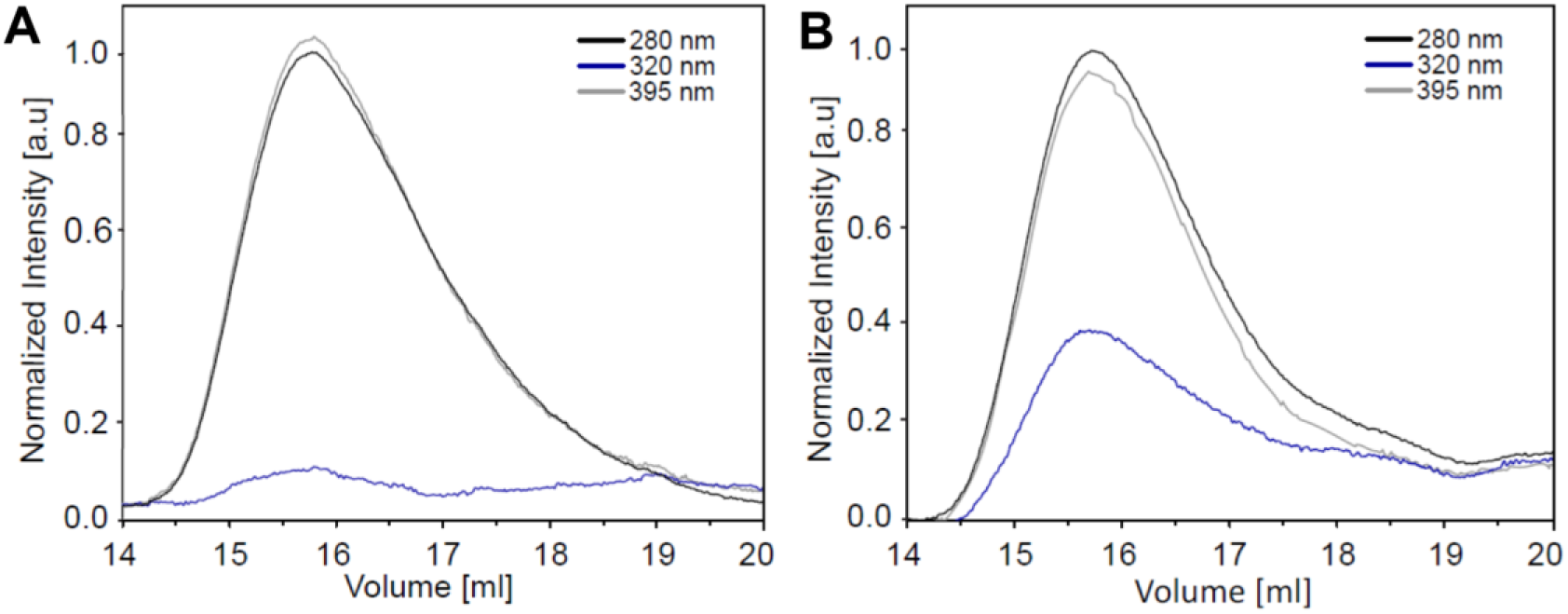
Size exclusion chromatograms of GFP-QC2 without **(A)** and with **(B)** 4-PAM showing an absorbance increase at 320 nm where PAM shows its maximum absorbance.

The procedure was repeated for the other three photostabilizers, although labelling could not be monitored by UV/VIS methods, because NPP, TX and COT show no characteristic absorbance at wavelengths >300 nm. Therefore, for these GFP-photostabilizer conjugates (GFP-COT, GFP-NPP, and GFP-TX), their spectroscopic characterization was performed using single-molecule TIRF (total internal reflection fluorescence) microscopy. The bulk emission spectra of unlabeled and all four labeled GFP-QC2 proteins were indistinguishable (Figure 2D) supporting the idea that no static complexes between photostabilizer and chromophore were formed, e.g., complexes with blue-shifted absorption spectra^27, 47^.

For single-molecule TIRF studies the proteins were immobilized on microscope coverslips according to published procedures^34^ (details see Material and methods). Unlabeled GFP-QC2 fluorophores were observed as well-separated diffraction-limited fluorescence spots in camera images (Figure 4A).

**Figure 4.**
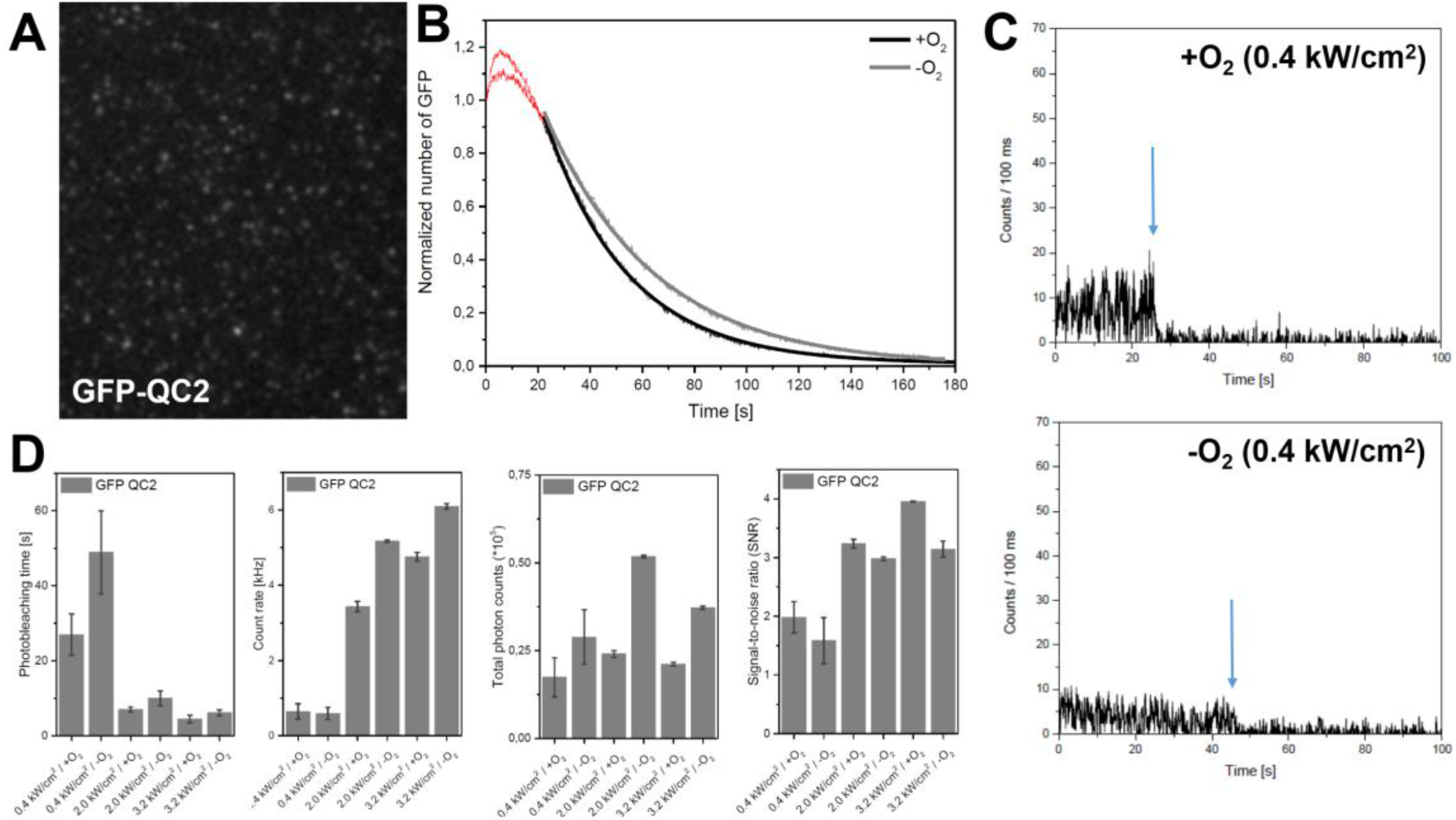
Quantitative photophysical characterization of GFP-QC2 in the presence and absence of oxygen under different excitation conditions following methods described in ref. ^34^. **(A)** TIRF image with **(B)** bleaching analysis counting fluorophore number per frame as a function of time. **(C)** Fluorescent time traces of individual GFP-QC2 molecules (arrows indicate photobleaching) with **(D)** quantitative photophysical analysis under different excitation conditions. All experiments were repeated within independent biological repeats for at least three tiems. Bar graphs were derived from averages of >5 movies per conditions per repeat.

GFP-QC2 behaved similarly to other fluorescent proteins when studied on the single-molecule level featuring low photostability (Figure 4B), poor signal-to-noise ratio (SNR) and low brightness for both oxygenated and deoxygenated conditions (Figure 4C). Deoxygenated conditions can increase photon emission as oxygen is a fluorescence quencher or diminish them if reactive-oxygen mediates novel photobleaching pathways ^47, 61–62^. The analysis of spot numbers in each movie frame (Figure 4B) and fluorescence time trace analysis (Figure 4C/5) using previously published procedures^34^ allowed us to quantitatively determine the count-rate, SNR and photobleaching times for single molecules for different excitation intensities (0.4, 2.0, 3.2 kW/cm^2^) in the absence and presence of oxygen (Figure 4D). For unlabeled GFP-QC2 fluorophores (Figure 4D), we observed short fluorescence periods of ~20 s with count rates of ~0.5 kHz at 0.4 kW/cm^2^ (see Figure 5 for individual traces). The SNR of GFP-QC2 at 100 ms binning was between 1.5-4 (Figure 4D).

**Figure 5.**
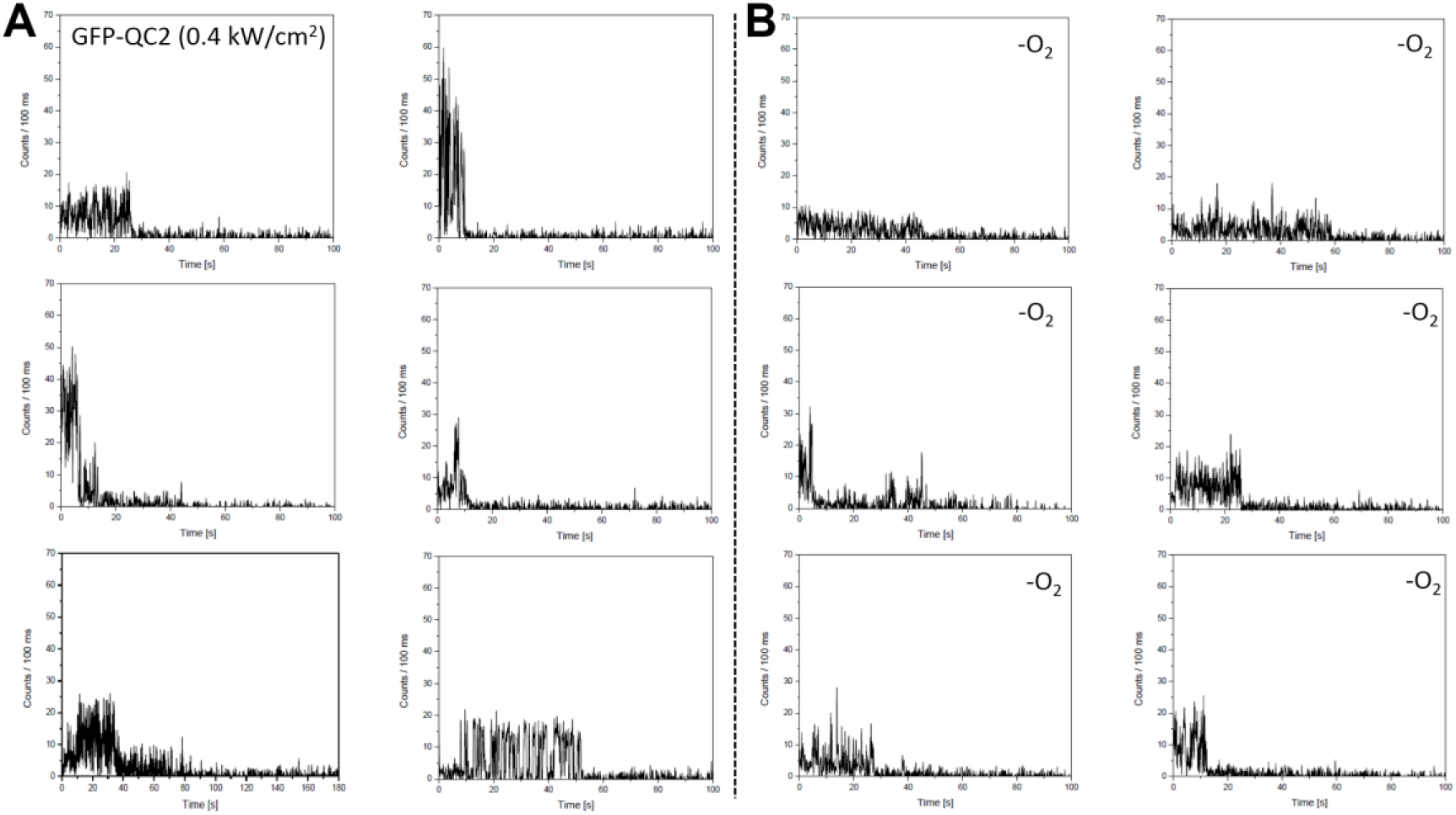
TIRF time traces of GFP-QC2 **(A)** in the presence and **(B)** in the absence of oxygen at 0.4 kW/cm^2^ excitation intensity.

The total number of detected photons were similar for most excitation conditions, i.e., between ~25000-50000. The constant values resulted from faster photobleaching but higher count-rate for increasing excitation intensity (Figure 4D). The normalized number of GFP-QC2 proteins per frame always showed an initial increase in the first 5-10 s that is consistent with previous reports of GFP/α-GFP and relates to photoconversion proesses (Figure 4B and ref. ^55^). We thus analyzed photobleaching times via an exponential fit of the tail of the decay. We also studied the influence of known solution additives such as COT and TX as controls (Figure S2). These experiments were done before we started our study on the intramolecular stabilizers to verify previous reports^51–53^ that solution additives (and thus potentially also molecules attached outside the ß-barrel) can influence the GFP-chromophore. For addition of both TX and COT, we found negative impacts on photobleaching rates, increased count-rate and constant total detected photons/SNR for single-immobilized GFP-QC2 molecules (Figure S2). Following these investigations, we tested covalent linkage of photostabilizers to the residues A206C and L221C (Figure 6).

**Figure 6.**
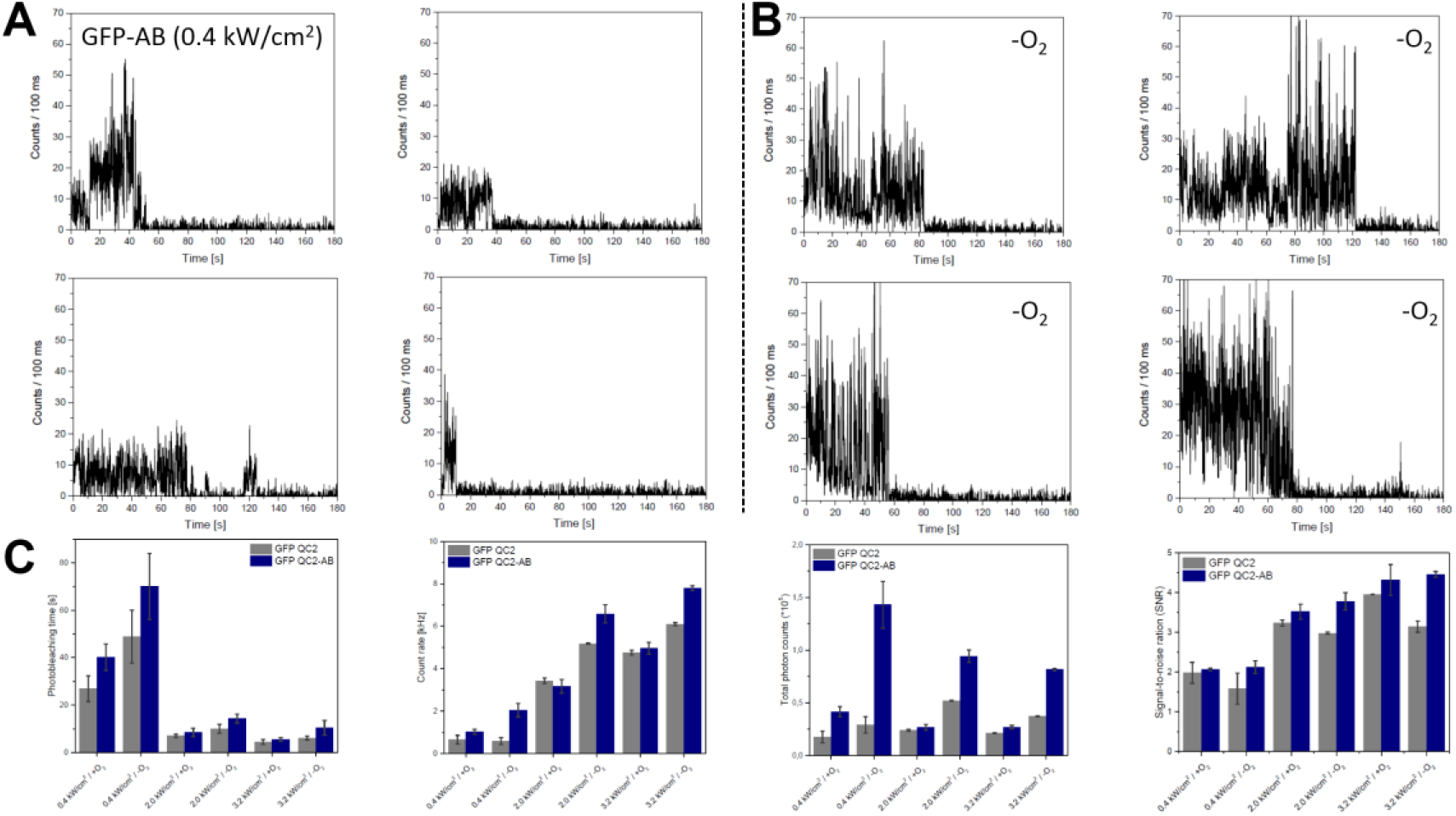
TIRF time traces of GFP-AB **(A)** in the presence and **(B)** in the absence of oxygen at 0.4 kW/cm^2^ excitation intensity. **(C)** Quantitative photophysical analysis of GFP-AB under different excitation conditions.

The selected photophysical parameters were improved by conjugation of 4-PAM to GFP-QC2, referred to as GFP-AB (Figure 6). Photobleaching was retarded by 4-PAM for all conditions (Figure 6C), but most significantly in the absence of oxygen. Increases in the count-rate by AB were only observed in the absence of oxygen. SNR changes were found to be non-systematic. Strikingly, the increases of both count-rate and photobleaching time gave rise to a substantial gain in the total number of observed photons before photobleaching for all excitation conditions, especially in the absence of oxygen (Figure 6C).

As outlined before, the barrel of GFP-QC2 was also labeled with the photostabilizers TX, NPP, and COT to generate GFP-TX, GFP-NPA, GFP-COT, respectively (Figure 7); see SI for synthesis of photostabilizer maleimides and the labelling procedure.

**Figure 7:**
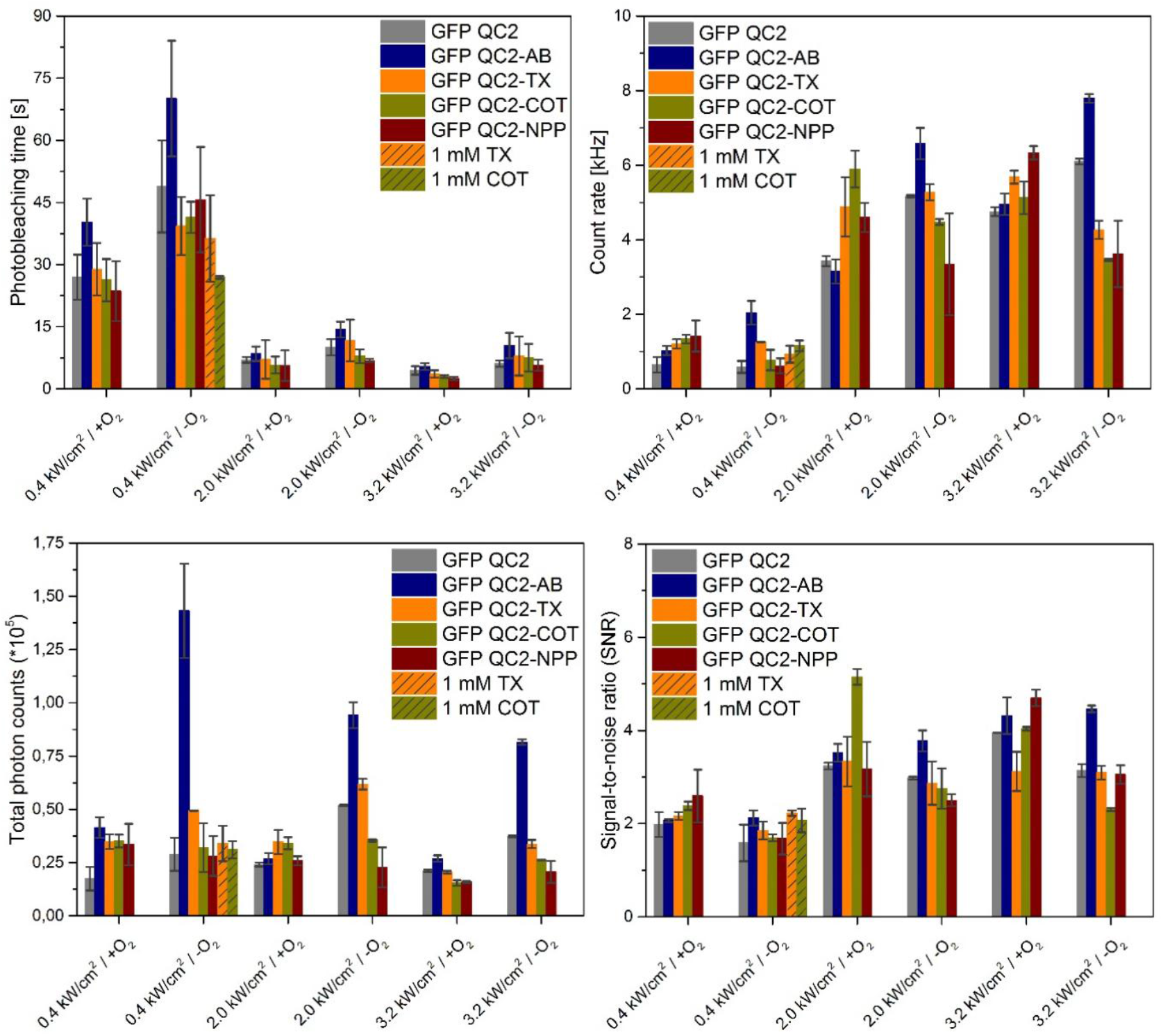
Quantitative photophysical characterization of GFP-QC2 with and without different photostabilizers in the presence and absence of oxygen at under different excitation conditions.

These experiments revealed only minor effects of the different stabilizers on the photophysical behavior of GFP-QC2 in contrast to 4-PAM. None of these other photostabilizers increased or decreased the photobleaching time, count-rate, total photon count and SNR strongly. Trolox showed some exceptions of this general statement with elevated count-rates at 2 kW/cm^2^.

The observed small effects of TX, NPP, and COT were on one hand disappointing, albeit not surprising since other blue fluorophores (Cy2^22^, fluoresceins^36^) were shown to be only minimally affected by these stabilizers. Importantly, these data further support that the idea of a unique photophysical interaction between the FP-chromophore and 4-PAM, which was not seen with any other stabilizer.

## 3. Summary and Discussion

In this study, we showed that a mutant GFP with two specific cysteine (A206/L221C) residues available for labelling with commercial and custom-made maleimide-photostabilizers, exhibited increased photostability upon conjugation to the azobenzene derivative 4-PAM (abbreviated GFP-AB). It could, however, not be shown that the underlying mechanism for this improvement is related to triplet-state quenching. Exactly this was demonstrated to be true for the class of self-healing dyes, which feature similar covalent linkage of photostabilizers to fluorophores^28^. The observed positive impact of 4-PAM on GFP photostability and the long recently determined triplet-state lifetimes of FPs^39^, however, supports the idea that FPs may be usefully targeted by intramolecular photostabilization, which provides an alternative approach to previous FP-improvement strategies using e.g., chromophore fluorination^63^.

While our study paves the way for a systematic investigations of how to equip GFPs with suitable intramolecular photostabilizers, there are several issues that require further attention. The strategy to label GFP on the outside of the ß-barrel may reduce efficient interaction between the chromophore and the photostabilizer. While, there is convincing published evidence that the ß-barrel does not shield the FP-chromophore fully^51–53^ from interacting molecules in the buffer and also that triplet-quenching proesses might be mediated by a contact-less Förster mechanisms^50^, we speculate that selecting a residue inside the ß-barrel might be even more promising. This could be done with residues such as C70 or other selected positions. In this case, a modified labelling strategy would be required, where the GFP is immobilized for labelling, unfolded to make the internal residue accessible and refolded after labelling has occurred.

Ultimately, a major point of discussion is the type of photostabilizer and quenching mechanism (PET vs. energy transfer) required to successfully stabilize GFP. As for a number of blue-absorbing fluorophores (Cy2 or fluorescein), the common quenchers TX, NPP and COT were also ineffective for GFP. Fluorescein and other blue dyes have a triplet energy of 1.98 eV, which is much higher than those found for green- and red-emitting dyes with values between 1.46 eV (ATTO647N) and 1.72 eV (TMR)^36^. The triplet-state of GFP was recently characterized and found to have a surprisingly low energy in the range of ~1.4 eV.^39^ This finding is not fully consistent with the fact that COT remains ineffective for GFP-QC2, since COT is very effective for ATTO647N, which has a similar triplet-state energy as GFP. Generally, for blue fluorophores alternative quenchers with energetically higher-lying triplet-states such azobenzene (~2 eV^56^), stilbene (~2.4 eV^64^) might be more optimal, also as solution additive for dyes with absorbance in the near-UV and blue spectral range.

## Acknowledgment

This work was financed by an ERC Starting Grant (No. 638536 – SM-IMPORT to T.C.) and Deutsche Forschungsgemeinschaft (SFB863 project A13 & GRK2062 project C03 to T.C. and JU650/2-2 to G.J.). L. Zhang thanks the Alexander von Humboldt foundation for a postdoctoral research fellowship. J.H.M.vdV. acknowledges Ubbo-Emmius funding (University of Groningen). T.C. was further supported by Deutsche Forschungsgemeinschaft through the cluster of excellence CiPSM and by the Center of Nanoscience Munich (CeNS). We thank D. A. Griffith for sequencing of the GFP-QC2 plasmid, reading of the manuscript and thoughtful comments and suggestions. We thank J. H. Smit and S. Franz for support and discussions in the initial phase of the project.

## Supplementary information

### 1. Additional data and images

**Figure S1:**
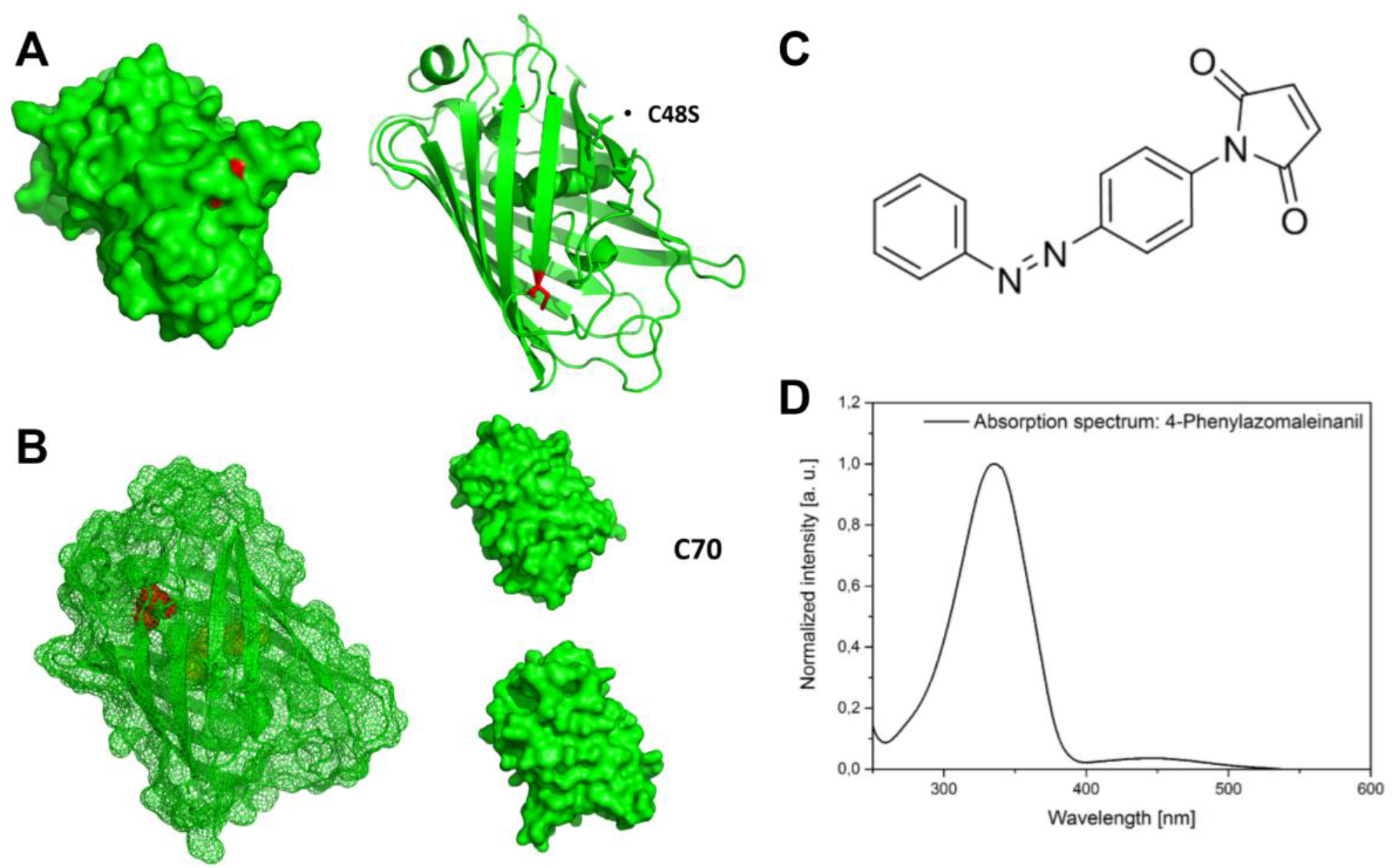
Crystal structures of GFP marking the location of **(A)** serine 48 (point mutation C48S, red) and **(B)** cysteine 70 (red). C48S is too far away from the chromophore and was thus deleted while C70 is not solvent-accessible in the folded form of GFP rendering both poor candidates for labelling of GFP with photostabilizers in the folded form of the protein. **(C)** Absorbance spectrum **(D)** and chemical structure of 4-phenylazomaleinanil (4-PAM) used for labelling of cysteine residues.

**Figure S2:**
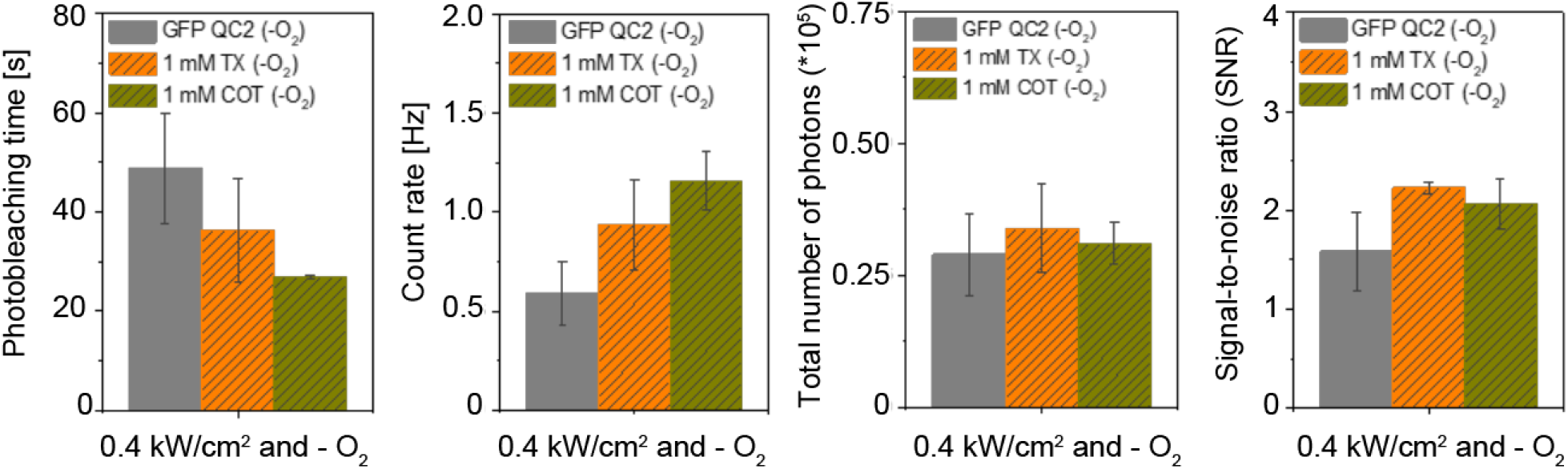
Photophysical properties of GFP-QC2 in different buffer environments in the absence of oxygen: no photostabilizer (grey), 1 mM TX (yellow) and 1 mM COT (green).

**Figure S3:**
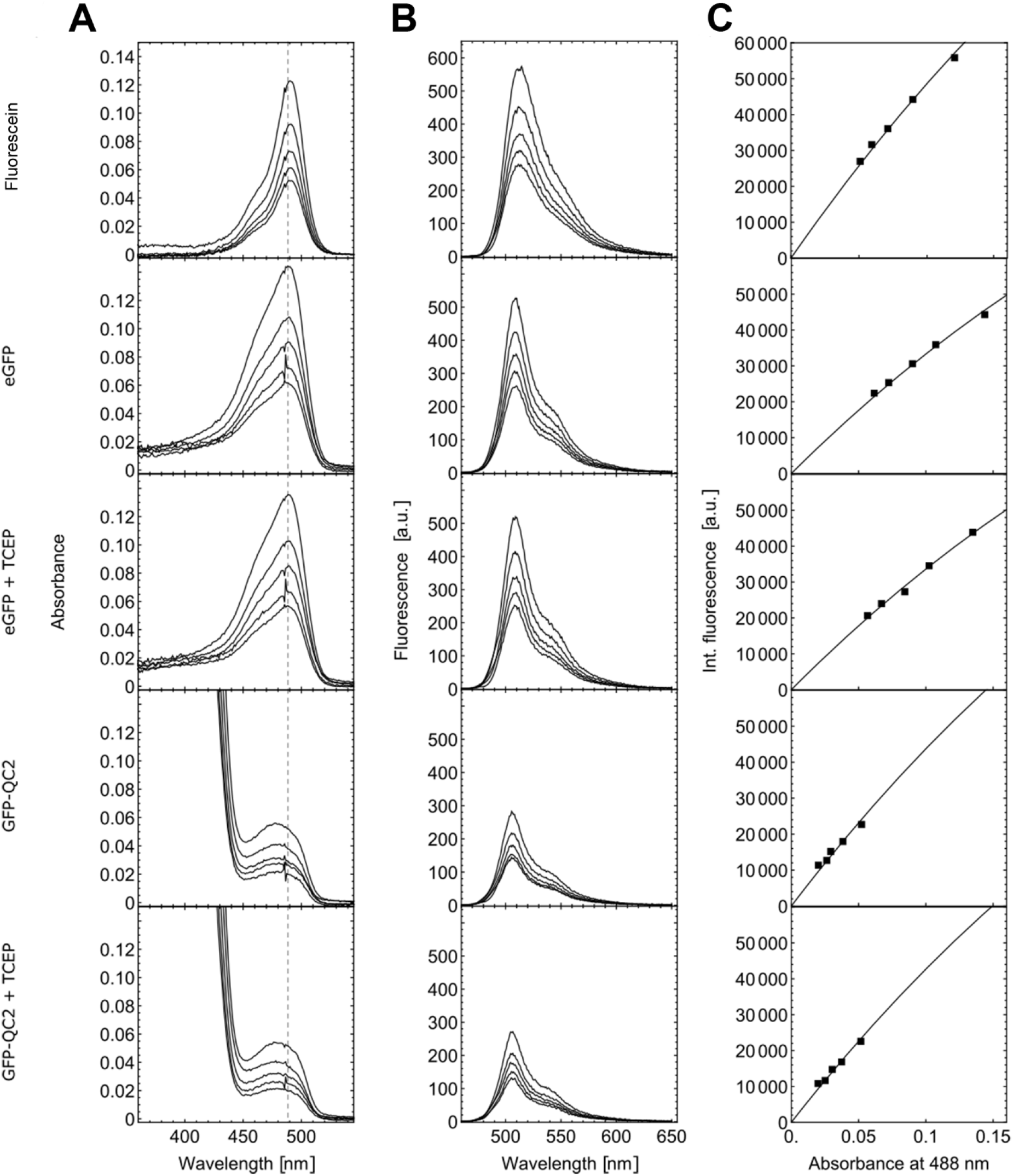
Quantum yield determination of eGFP and GFP-QC2 using fluorescein as standard. **(A)** Absorbance spectra with marked line at 488 nm, **(B)** emission spectra from excitation at 488 nm, and **(C)** integrated emission spectrum from (B) versus the absorbance at 488 nm from (A) with fitted curve 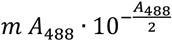 for Fluorescein, eGFP (without and with 1mM TCEP), and GFP-QC2 (without and with 1mM TCEP) (top to bottom). All measurements were done at 5 different concentrations. eGFP at 0.67, 0.50, 0.40, 0.33 and 0.29 mg mL^−1^ concentration, GFP-QC2 at 0.93, 0.69, 0.56, 0.46 and 0.40 mg mL^−1^ concentration, and fluorescein at 1.75, 1.31, 1.04, 0.87, 0.74 μM concentration.

### 2. Material and Methods

For all methods described below, chemicals and conjugates from the companies Sigma-Aldrich, Merck KGaA, Roche Diagnostics GmbH, J. T. Baker, abcr GmbH, Laysan Bio, Qiagen and Macron Fine Chemicals were used without further purification.

#### Overexpression and purification of GFP-QC2

The GFP variant used here, as a starting point for the construction of GFP-QC2, was the Stemmer cycle 3 mutant or αGFP (F99S/M153T/V163A)^1^. The αGFP gene was subcloned in frame with a hexa-histidine tag sequence to produce a C-terminal His_6_ fusion protein. The C48S, A206C, and L221C mutations were introduced by Quick-Change site-directed mutagenesis to produce the final plasmid pGFP-QC2 (see Figure S4 for plasmid map). The sequence of the GFP-QC2 gene was verified by di-deoxy sequencing. The plasmid was used to transform the *E. coli* BL21(DE3) strain (New England Biolabs). For protein expression, a single colony of *E. coli* BL21(DE3) carrying the expression construct was selected and grown in LB medium supplemented with 100 μg/mL ampicillin at 37°C overnight. The next day, overnight culture was used to inoculate 1 L of LB containing 100 μg/mL ampicillin. At an optical density (OD_600 nm_) of 0.6-0.8, expression of the GFP-QC2 cysteine mutant was induced by adding IPTG to 1 mM and growing for 3-4 h at 30°C. Following centrifugation, the cell pellet was resuspended and stored in 50 mM Tris, 1M KCl, 1% (v/v) glycerol, 1 mM DTT, 5 mM imidazole (pH 8.0) at −20°C.

**Figure S4:**
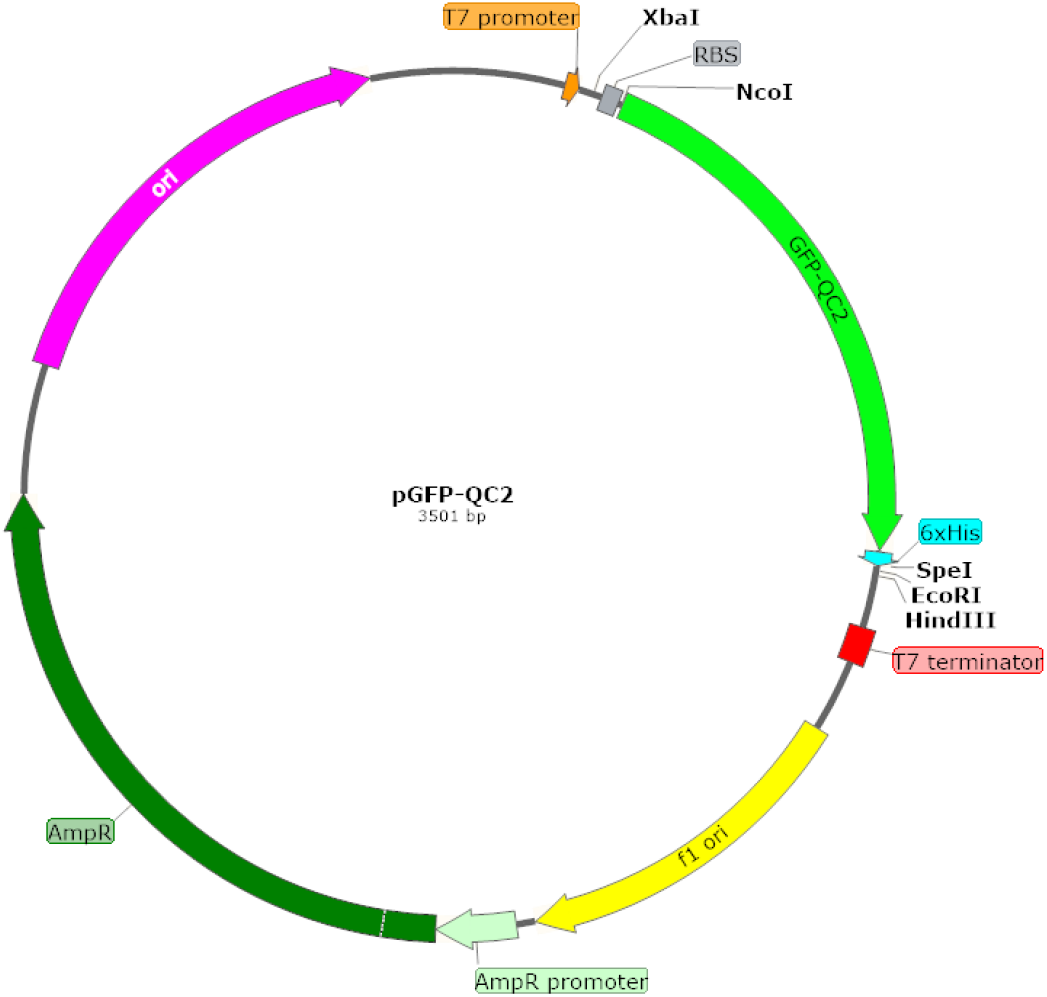
Physical and functional map of pGFP-QC2 plasmid. Relevant features of pGFP-QC2 are annotated on the map in different colours: T7 promoter (orange), ribosome binding site (RBS, gray box), GFP-QC2 gene (green) with C-terminal His^6^-tag (cyan), T7 terminator (red box), F1 origin (yellow), ampicillin resistance gene (AmpR) promoter (pale green), AmpR (dark green), and ColE1-like origin of replication (magenta). Unique restriction sites around GFP-QC2 are indicated. All genes are reported in scale over the total length of the vector. Images were obtained by the use of SnapGene software (from GSL Biotech).

Before cell lysis, if necessary, cell pellets were resuspended in 50 mM Tris, 1M KCl, 1 mM DTT, 5 mM imidazole (pH 8.0). Cell lysis was performed by adding lysis buffer (50-100 μg/mL DNAse, 1 mM MgCl_2_ and 1 mM DTT) followed by mechanical cell disruption using TissueLyser LT (Qiagen). After complete cell lysis, ethylenediaminetetraacetic acid (EDTA) and phenylmethylsulfonyl fluoride (PMSF) were added to final concentration of 5 mM (pH 7.4) respectively 1 mM. Clarified extract was collected following centrifugation at 40k rpm for 1 h at 4°C (Beckman Coulter, Avanti J-20 XP Centrifuge).

His_6_-tagged GFP-QC2 cysteine mutant was purified from clarified extract by nickel-affinity chromatography. First, nickel resin was washed with ten volumes ethanol, MilliQ water and equilibrated with ten column volumes of Equilibration Buffer (Table S1). Clarified extract was then loaded on column followed by washing with ten column volumes of Washing Buffer (Table S1). His_6_-tagged GFP-QC2 cysteine mutant was then eluted from nickel column using Elution Buffer (Table S1). To evaluate purification progress, reduced samples of supernatant, flow through, wash steps and the elution steps were loaded onto SDS-PAGE gel (Figure S5).

**Figure S5:**
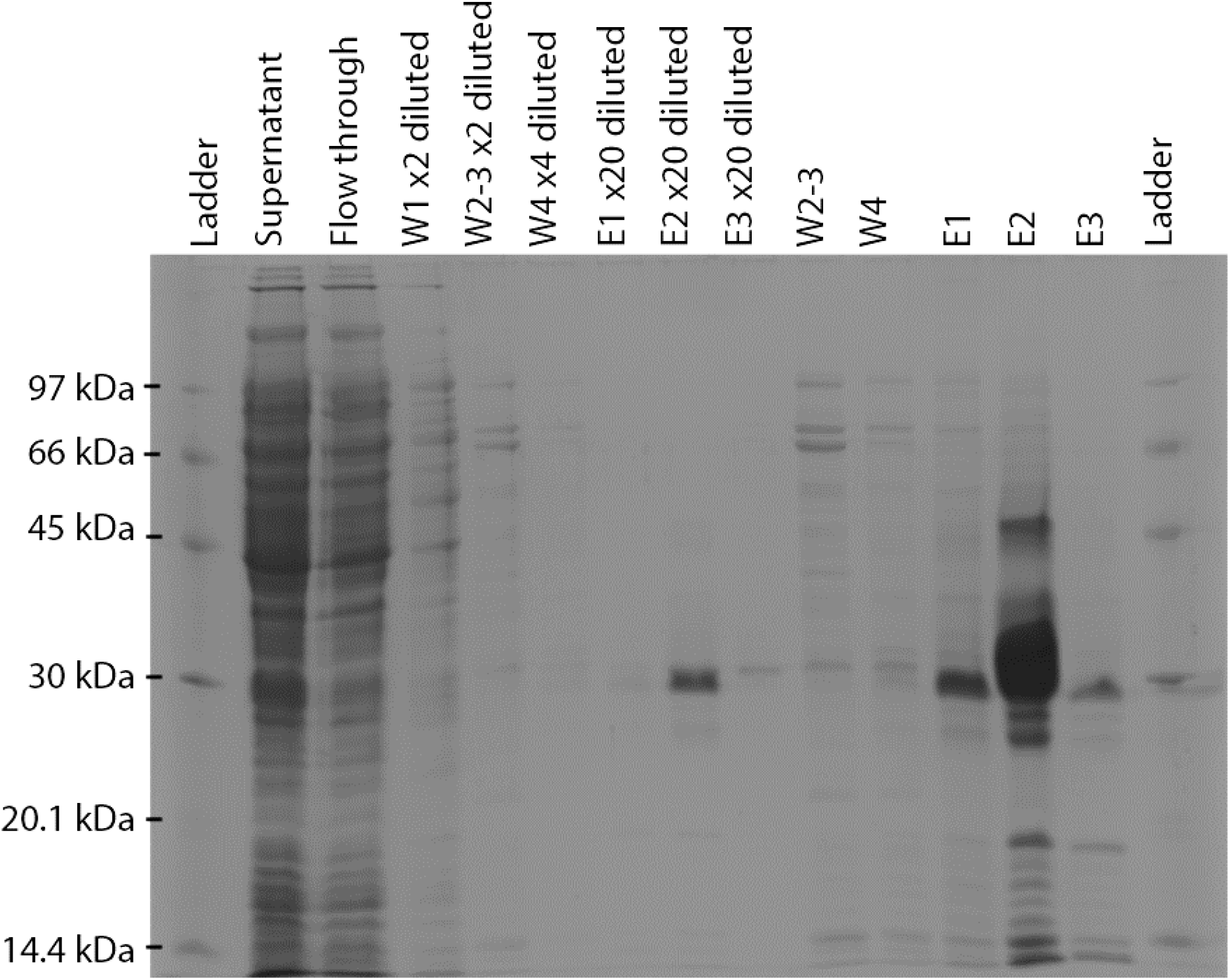
SDS-PAGE gel showing purification steps of GFP-QC2 using nickel-affinity column. Lanes 1: low molecular ladder (LMW-SDS Marker Kit, GE Healthcare Europe GmbH); 2: supernatant; 3: flow through; 4: wash 1 diluted by a factor of two; 5: wash 2-3 diluted by a factor of two; 6: wash 4 diluted by a factor of four; 7: elution 1 diluted by a factor of 20; 8: elution 2 diluted by a factor of 20; 9: elution 3 diluted by a factor of 20; 10: wash 2-3 undiluted; 11: wash 4 undiluted; 12: elution 1 undiluted; 13: elution 2 undiluted; 14: elution 3 undiluted; 15: low molecular ladder. SDS-PAGE gel was run in two intervals: 1. 10 min at 100 V and 2. 60-90 min at 200 V.

Protein eluted from nickel column was concentrated by ultrafiltration (Amicon Ultra 4, 10,000 molecular weight cut-off (MWCO), Merck KGaA). Using concentrated protein in dialysis system (SnakeSkin™ Dialysis Tubing, 10K MWCO, 22 mm, Thermo Fisher Scientific), buffer was exchanged to storage buffer (50 mM Tris-HCl pH 8, 50 mM KCl, 50% (v/v) glycerol, 1 mM DTT). Dialysis was performed in two stages at 4 °C, with ≥ 12 h for each dialysis stage. Buffers for dialysis stage 1 and stage 2 are listed in Table S1. Following dialysis, 3 mM EDTA (pH 7.4) and 1 mM DTT were added and protein stock was stored at −80 °C. Protein concentration was determined by the bicinchoninic acid method (PierceTM BCA Assay Kit, Thermo Fisher Scientific) with bovine serum albumin as the standard and absorption measurements (NanoDrop ND-1000 Spectrophotometer, NanoDrop Technologies).

#### Labelling of GFP-QC2 with photostabilizers

GFP-QC2 cysteine was modified in a reaction with a photostabilizer-maleimide derivatives (AB-Mal, TX-Mal, NPP-Mal or COT-Mal)^2–3^, coupling GFP-QC2 cysteine with the maleimide group. Briefly, cysteines were first reduced by adding 5 μL of 425 μM GFP-QC2 (2.1 nmol) to 95 uL of DTT-containing buffer (50 mM potassium phosphate buffer [KPi buffer], 50 mM KCl, 5% glycerol [v/v] pH 7.4, 5 mM DTT). Following 30 min incubation, protein solution was mixed with 1 mL standard buffer (50 mM potassium phosphate buffer [KPi buffer], 50 mM KCl, 5% glycerol [v/v] pH 7.4) and subsequently loaded on 150 μL nickel resin (Ni Sepharose, 6 Fast Flow, GE Healthcare Europe GmbH) equilibrated with 1 mL standard buffer. DTT was then washed off using ten column volumes of standard buffer. Maleimide-cysteine coupling was carried out on the resin by adding a solution of 1 mL standard buffer and 10 μL DMSO containing 100 nmol photostabilizer. The reaction was incubated overnight at 4°C with gentle shaking. The next day, the resin was washed with ten column volumes of standard buffer, before eluting the protein with 1 mL buffer containing 500 mM imidazole, 50 mM KPi, 50 mM KCl, 5% glycerol (v/v). GFP-QC2 photostabilizer conjugate was further purified by size exclusion chromatography, removing excess of unbound photostabilizer, which at the same time allowed us to assess the labelling efficiency. Labelling efficiency for 4-PAM was further determined by measuring absorbance increase at 320 nm (Figure 3, main text).

#### Sample preparation for single-molecule imaging

Lab-Tek 8-well 750 μL chambered cover slides (#1.0 Borosilicate Coverglass System, Nunc/VWR, The Netherlands) were cleaned by incubating with 0.1 M HF for 10 min and rinsing three times with PBS buffer (10 mM phosphate, 2.7 M KCl, 137 mM NaCl at pH 7.4, Sigma-Aldrich)^4^. After cleaning, an affinity surface was generated for his_6_-tagged GFP-QC2. First, cleaned cover slides were biotinylated by incubating with a solution of 3 mg/mL BSA (Roche Diagnostics GmbH) and 1 mg/mL BSA-biotin (Sigma-Aldrich) at 7 °C for 3-4 h. After rinsing with PBS, cover slides were incubated with 0.2 mg/mL streptavidin dissolved in PBS for 10 min at room temperature, binding streptavidin to biotinylated surface^5^. Non-bound streptavidin was washed off with PBS. Finally, each chamber was incubated with 1 μL Penta·His_6_ Biotin Conjugate (Qiagen) in 200 μL deionized water for 10 min and subsequently rinsed with PBS buffer. Derivatization steps resulted in free Penta·His_6_ groups on the surface (Figure S6), forming an affinity surface for his_6_-tagged protein.

Immobilisation of his_6_-tagged GFP-QC2 and photostabilizer-protein conjugates allows the characterization of photophysical properties. To homogeneously cover the glass surface, 20 μL of 5 nM GFP sample in 200 μL MilliQ water were added to a chamber which was subsequently rinsed with a high concentrated salt solution (1 M KPi) and PBS^4^. If applicable, buffer was deoxygenated in chambers^4^ by using an oxygen scavenging system (PBS buffer at pH 7.4 including 1% (w/v) glucose and 10% (w/v) glycerol, 50 μg/mL glucose oxidase, 100-200 μg/mL catalase, 0.1 mM tris(2-carboxyethyl)phosphine hydrochloride [TCEP]) for which the chambers were sealed with adhesive tape (Adhesive silicon sheet JTR-SA2-2.5, Grace Bio-Labs).

**Figure S6:**
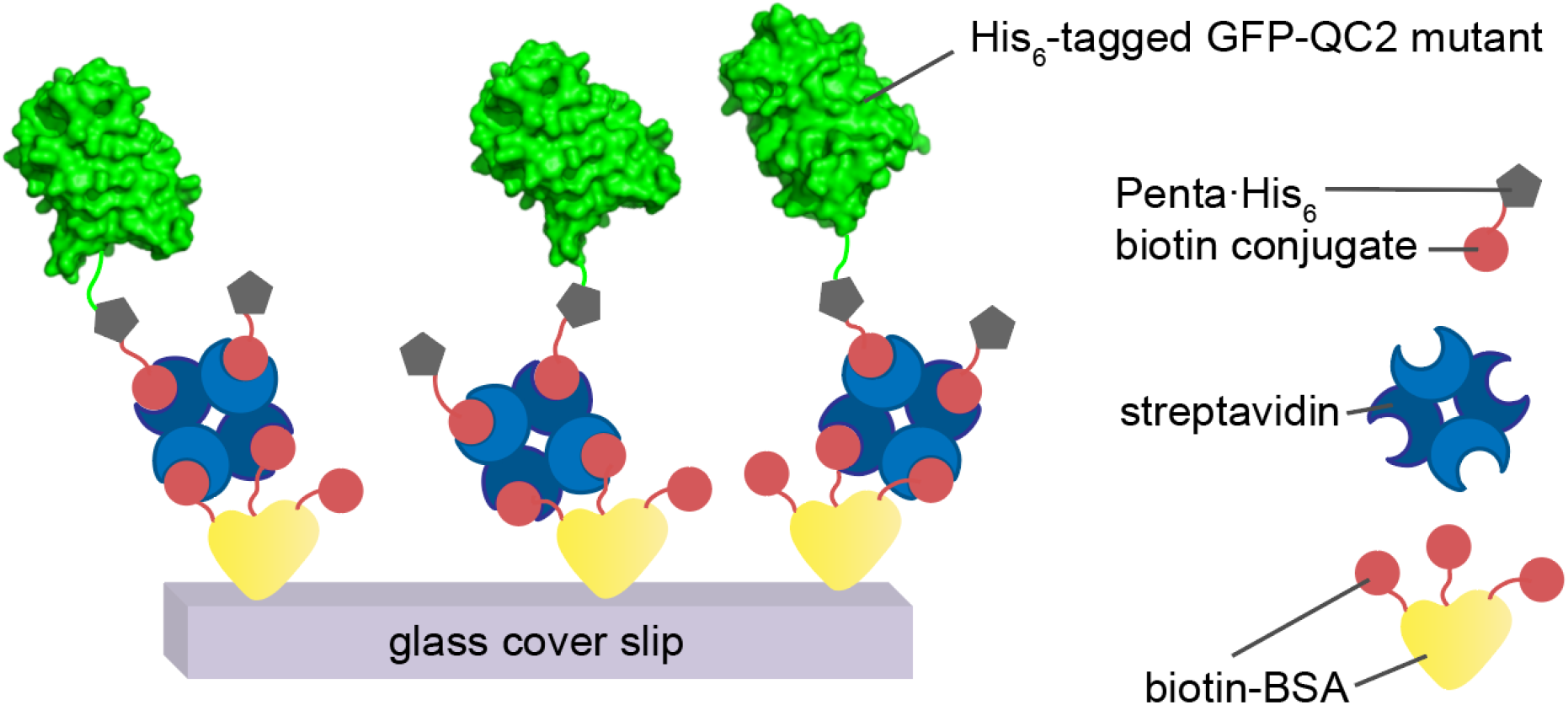
Immobilisation of GFP-QC2 on a affinity-surface, prepared on Lab-Tek coverglass system.

#### Spectroscopy & Quantum yield determination

Absorbance spectra were recorded using absorption spectrometer V-630 (wavelength accuracy ± 0.2 nm, photometric accuracy ± 0.002 Abs. [0 to 0.5 Abs.] and ± 0.002 Abs. [0.5 to 1 Abs.], JASCO) and quartz glass cuvettes (precision cuvettes made of quartz glass Model FP-1004, d = 1 cm, JASCO parts center). Fluorescence spectra were recorded with the fluorescence spectrometer FP-8300 (wavelength accuracy ± 1.5 nm, JASCO) and quartz glass cuvettes (precision cuvettes made of quartz glass Model FP-1004, d = 1 cm, JASCO parts center).

Fluorescence quantum yields were determined for eGFP and GFP-QC2 with 1 mM and without TCEP in PBS in comparison to the quantum yield standard fluorescein in 0.1 M NaOH^6^. The absorbance spectra and emission spectra obtained via 488 nm excitation were recorded for five different fluorophore/protein concentrations. Absorbance spectra were base-line corrected to remove buffer background. Emission spectra were corrected for wavelength-dependent detection efficiency and excitation scattering light. The integrated fluorescence 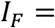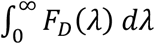 was obtained by recording the emission spectra *F*_D_(λ) introducing corrections for reabsorbance of the fluorescence. We estimated this via 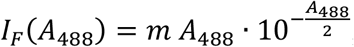, where the factor 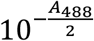 accounts for the absorption of excitation light during emission measurements.

The absolute fluorescence quantum yield of the GFP proteins (eGFP, GFP-QC2) were calculated from the slopes of the fits of GFP *m*_*GFP*_ and fluorescein *m*_*flcn*_ as

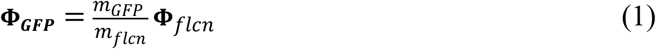

We obtained Φ_*flcn*_ = 92.5% from the literature^6^. The reported values and standard deviations resulted from three independent experiments.

#### Single-molecule TIRF imaging

Widefield fluorescence and TIRF imaging was performed on an inverted microscope (Olympus IX-71 with UPlanSApo 100x, NA 1.49, Olympus, Germany) in an objective type total-internal-reflection fluorescence (TIRF) configuration. The images were collected with a back-illuminated emCCD camera (512×512 pixel, C9100-13, Hammamatsu, Japan in combination with ET535/70, AHF Analysentechnik, Germany). Excitation is conducted from a diode laser (Sapphire and Cube, Coherent, Germany) at 488 nm with ≈ 0.4-3.2 kW/cm^2^ at the sample location. The imaging area covers a size of ≈ 25×35 μm containing >40 proteins and the full chip amounts to 50×50 μm. The recorded movies range over 100-180 s with an integration time of either 50 ms or 100 ms. Fluorescence time traces were extracted from pixels which showed at least 2-3 standard deviations above background noise (standard deviation of all pixels over all frames of the movie) and summing the intensity in a 3×3 pixel area. Neighbouring peaks closer than 5 pixels were not taken into account. The number of fluorescent spots in each frame image was determined using an absolute threshold criterion. The number of proteins per image are plotted over time [s] and fitted to a mono-exponential decay y(t)=C+A∙e^(−bt) (with b = 1/_τbleach_ and τ_bleach_ being the characteristic bleaching time constant). Using these fluorescent time traces, four photophysical properties were measured: 1.) Bleaching times and corresponding standard deviations were derived from multiple repeats of the same measurement on different days, where each condition was tested ≥ 2 movies. 2.) Signal-to-noise (SNR) ratio was determined by dividing the standard deviation of the signal before photobleaching with the average fluorescence intensity during that period. 3.) Count rate, respectively brightness, was obtained by multiplying the signal (counts / 100 ms / pixel) by 10 to receive counts / s / pixel, by 9 to gain counts / s and by 111.14 to obtain photons / s (conversion from counts to photons is a device-specific value for CCD camera). 4.) Total number of detected photons before bleaching were calculated by multiplying the count rate by τ_bleach_.

**Table S1:** Buffers and solutions and their final concentrations.

| **Buffer** | **Composition** |
| --- | --- |
| Equilibration Buffer | 50 mM Tris-HCl, pH 8<br>1 M KCl<br>1% (v/v) glycerol<br>1 mM DTT<br>5 mM imidazole |
| Wash Buffer | 50 mM Tris-HCl, pH 8<br>100 mM KCl<br>2% (v/v) glycerol<br>1 mM DTT<br>40 mM imidazole |
| Elution Buffer | 50 mM Tris-HCl, pH 8<br>100 mM KCl<br>2% (v/v) glycerol<br>10 mM DTT<br>300 mM imidazole |
| Dialysis 1 | 50 mM Tris-HCl, pH 8<br>50 mM KCl<br>5% (v/v) glycerol<br>1 mM DTT |
| Dialysis 2 | 50 mM Tris-HCl, pH 8<br>50 mM KCl<br>50% (v/v) glycerol<br>1 mM DTT |
| Stacking Buffer | 0.5 M Tris-HCl, pH 6.6<br>10% (w/v) sodium dodecyl sulfate (SDS) |
| Separation Buffer | 4.5 M Tris-HCl, pH 8.8<br>10% (w/v) SDS |
| SDS-Loading Buffer | 250 mM Tris-HCl, pH 6.6<br>10% (w/v) SDS<br>0.2% (w/v) bromophenol blue<br>12.5% (w/v) β-mercaptoethanol<br>50% (v/v) glycerol |
| SDS-Running Buffer | 250 mM Tris<br>1.92 M glycine<br>1 % (w/v) SDS |
| Coomassie Brilliant Blue | 0.125% (w/v) Coomassie R<br>40% (v/v) ethanol<br>5% (v/v) acetic acid |

### 3. Synthesis and characterization of photostabilizer-maleimide derivatives

#### AB-Mal

4-phenylazomaleinanil (4-PAM, see Figure S1) was purchased from Sigma Aldrich (CAS Number 103-33-3) with 98% purity.

#### NPP-Mal

NPP-Mal was obtained by coupling 3-(4-nitrophenyl)propanoic acid (NPP) with 1-(2-aminoethyl)-1*H*-pyrrole-2,5-dione (maleimide amine, Mal-NH_2_) following a modified procedure^7^ (see Figure S7). Briefly, NPP (1.0 equiv, 20.1 mg, 0.1 mmol) and Mal-NH2 (3.6 equiv, 93.3 mg, 0.37 mmol) were dissolved in 1 mL DMF and HATU (5.2 equiv, 0.21 g, 0.54 mmol) in 0.5 mL DMF was added.

**Figure S7:**
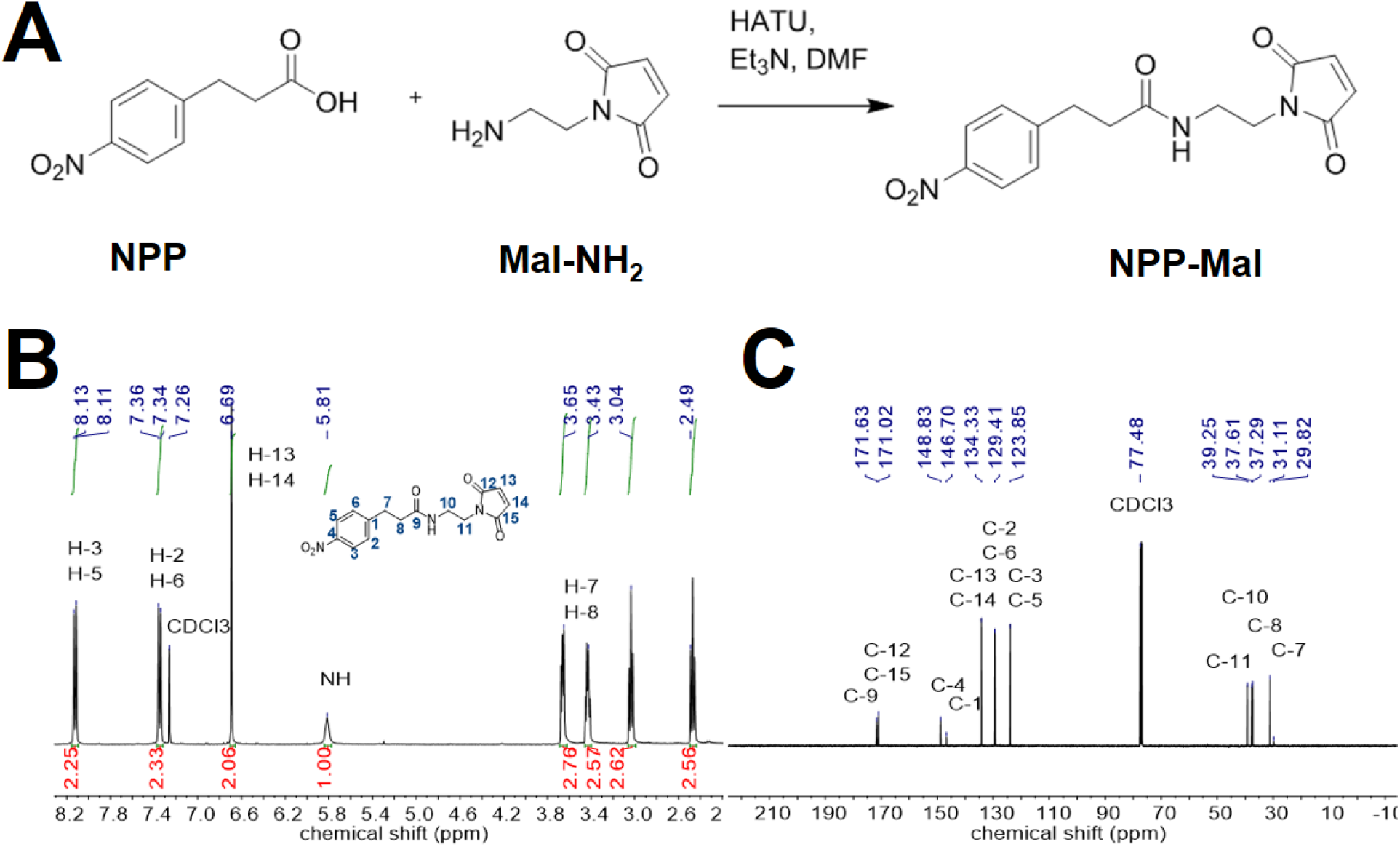
**(A)** Coupling scheme for synthesis of NPP-Mal. **(B)** ^1^H spectrum and **(C)** ^13^C spectrum of NPP-Mal.

Then, Et_3_N (50 μL) was added dropwise to the solution and the reaction mixture was stirred at room temperature for 19.5 h. The reaction mixture was concentrated and the crude product was purified by column chromatography (SiO2, DCM/MeOH 99:1) yielding a yellowish solid (30.3 mg, 0.09 mmol, 93 %). The product was characterized by NMR spectroscopy and mass spectrometry (see Figure S7).

^1^H NMR (400 MHz, CDCl3) δ = 8.13 (s, 1H, H-3), 8.11 (s, 1H, H-5), 7.36 (s, 1H, H-2), 7.34 (s, 1H, H-6), 6.69 (s, 2H, H-13, H-14), 5.81 (br s, 1H, NH), 3.65 (tr, *J* = 5.3 Hz, 2H, H-7), 3.43 (quart, *J* = 5.2 Hz, 2H, H-8), 3.04 (tr, *J* = 7.7 Hz, 2H, H-11), 2.49 (tr, *J* = 7.7 Hz, 2H, H-10) ppm.

^13^C NMR (200 MHz, CDCl3) δ = 171.63 (C-9), 171.02 (C-12, C-15), 148.83 (C-4), 146.70 (C-1), 134.33 (C-13, C-14), 129.41 (C-2, C-6), 123.85 (C-3, C-5), 39.25 (C-11), 37.61 (C-10), 37.29 (C-8), 31.11 (C-7) ppm.

Mass spectrometry (ESI, *full scan*) m/z calculated 317.29682, found 318.10846 [M+H]^+^, 340.09034 [M+Na]^+^, 356.06425 [M+K]^+^.

#### TX-Mal

TX-Mal was obtained in a two-step reaction. Frist, Trolox-NHS was synthesized following a modified procedures^2,8^. Trolox (TX) (1.0 equiv, 0.282 g, 1.13 mmol) and *N*-hydroxysuccinimide (NHS) (1.2 eqiv, 0.251 g, 1.33 mmol) were dissolved in 4.5 mL 1,4-dioxane. The reaction mixture was cooled to 0°C and *N,N’*-dicyclohexyl carbodiimide (DCC) (0.7 equiv, 0.155 g, 0.75 mmol) was added. The resulting mixture was allowed to warm up to room temperature and stirred for 19 h. Following reaction, the mixture was cooled to 10 °C, filtered and concentrated. To remove residue 1,4-dioxane, anhydrous ethanol was added and evaporated. The crude prodcut was purified by column chromatography (SiO2, DCM/MeOH 99:1) to produce a white solid of TX-NHS (65.2 mg, 0.19 mmol, 17%). The product was confirmed by NMR spectroscopy: ^1^H NMR (400 MHz, CDCl3) δ = 2.73 (s, 4H, H-16, H-17), 2.69-2.66 (m, 1H, H-3a), 2.58-2.53 (m, 1H, H-3b), 2.15 (s, 3H, H-11), 2.13 (s, 3H, H-12), 2.07 (s, 3H, H-10), 2.04-1.96 (m, 1H, H-2), 1.82 (s, 3H, H-13) ppm.

To generate TX-Mal, purified Trolox-NHS was coupled with 1-(2-aminoethyl)-1*H*-pyrrole-2,5-dione (Mal-NH_2_) following a published procedure^2^ (see Figure S8A). TX-Mal (1.0 equiv, 0.065 g, 0.19 mmol) was dissolved in 2.5 mL DMF and a solution of Mal-NH_2_ (1.5 equiv, 0.072 g, 0.51 mmol) and Et3N (50 μL) in 1.5 mL DMF was added. This mixture was stirred for 18 h at room temperature. At that point, 1 mL of water was added and the solution was acidified with H_2_SO_4_ to pH 1. The reaction mixture was extracted with EtOAc (3 × 5 mL), the combined organic phases were dried over Na2SO4 and concentrated. The crude product was purified by gradient column chromatography (SiO2, DCM/MeOH 99:1 – 95:5) to amount to a yellowish solid (2.8.9 mg, 0.08 mmol, 41 %). The product TX-Mal was confirmed by NMR spectroscopy and mass spectrometry.

^1^H NMR (400 MHz, CDCl3) δ = 8.01 (s, 1H, NH), 6.62 (s, 1H, OH), 6.57 (s, 2H, H-18, H-19), 3.72 – 3.37 (m, 4H, H-2, H-3), 2.95 (s, 2H, H-16), 2.88 (s, 2H, H-15), 2.16 (s, 6H, H-11, H-12), 2.07 (s, 3H, H-10), 1.45 (s, 3H, H-13) ppm.

^13^C NMR (400 MHz, CDCl3) δ = 175.09 (C-14), 170.66 (C-17, C-20), 145.63 (C-6), 144.26 (C-9), 133.95 (C-18, C-19), 122.18 (C-7), 121.45 (C-5), 119.02 (C-4), 118.09 (C-8), 78.36 (C-1), 37.97 (C-16), 37.40 (C-15), 29.51 (C-3), 24.53 (C-2), 20.55 (C-13), 12.38 (C-10), 12.10 (C-11), 11.46 (C-12) ppm.

Mass spectrometry (ESI, *full scan*) m/z calculated 372.41504, found 373.17508 [M+H]^+^, 395.15694 [M+Na]^+^, 411.15176 [M+K]^+^, 767.32478 [M2+Na]^+^.

**Figure S8:**
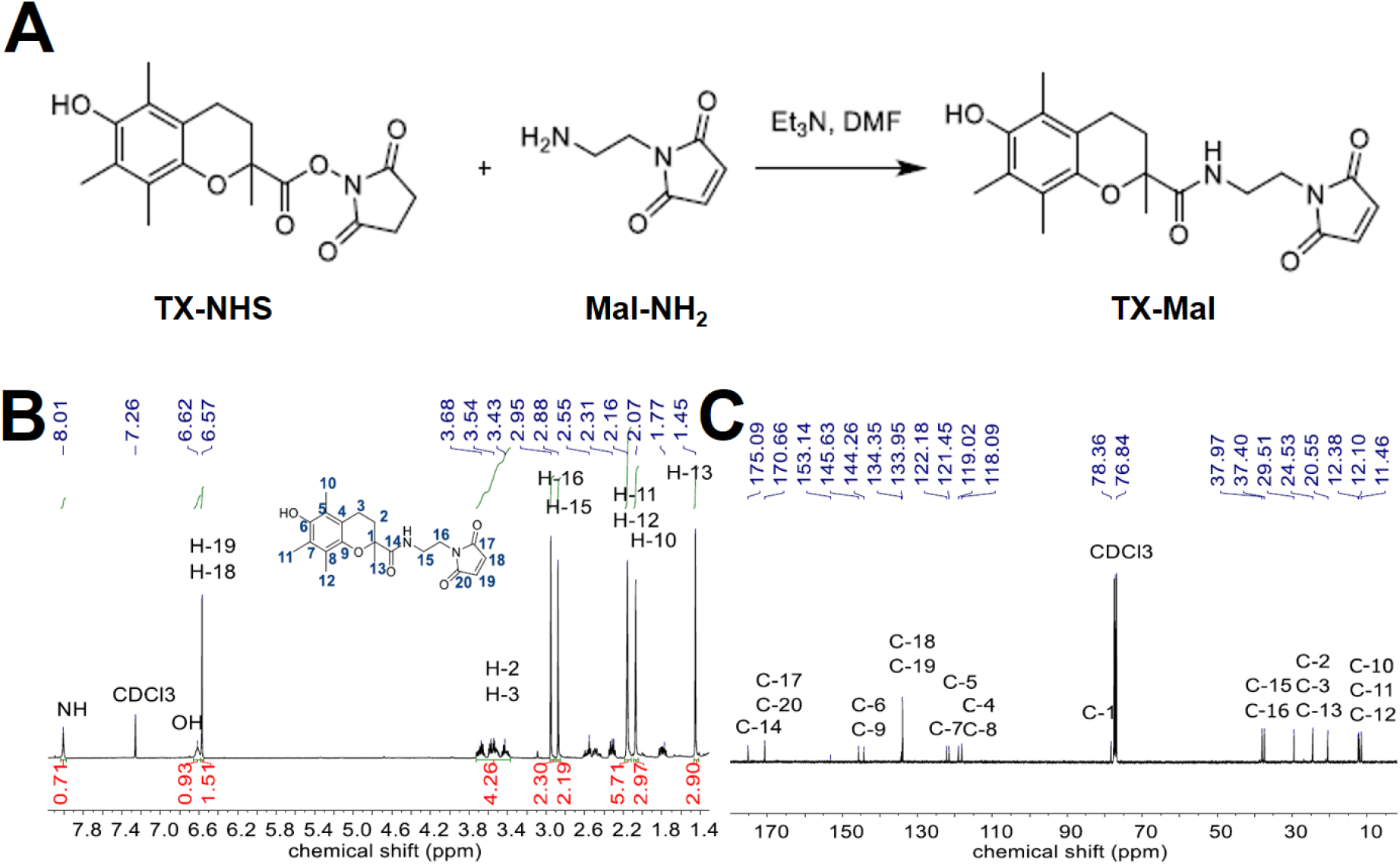
**(A)** Coupling scheme for synthesis of TX-Mal. **(B)** ^1^H spectrum and **(C)** ^13^C spectrum TX-Mal.

#### COT-Mal

COT-Mal was synthesized by forming an amide bond between COT-COOH and Mal-NH_2_ (see Figure S9A), following a modified published procedure^2^. Educt COT-COOH was previously synthesized^9^. Mal-NH_2_ (1.0 equiv, 30.2 mg, 0.12 mmol) and COT-COOH (1.1 equiv,22.1 mg, 0.13 mmol) were dissolved in 1 mL DMF and HATU (5.8 equiv, 0.26 g, 0.69 mmol) in 1.0 mL DMF was added. Then, Et3N (50 μL) was added dropwise to the solution and the reaction mixture was stirred at room temperature for 19.5 h. The reaction mixture was concentrated and the crude product was purified by column chromatography (SiO2, DCM/MeOH 98:2) to yield a yellowish solid (13.0 mg, 0.03 mmol, 25 %).

^1^H NMR (400 MHz, CDCl3) δ = 6.72 (s, 2 H, H-15, H-16), 5.92 (br s, 1H, NH), 5.88 – 5.67 (m, 6H, H-3, H-4, H-5, H-6, H-7, H-8), 5.60 (s, 1H, H-2), 3.69 (tr, *J* = 5.5 Hz, 2H, H-13), 3.46 (quart, *J* = 5.5 Hz, 2H, H-12), 2.37 – 2.28 (m, 2H, H-10), 2.28 - 2.21 (m, 2H, H-9) ppm.

^13^C NMR (400 MHz, CDCl3) δ = 172.84 (C-11), 171.01 (C-14, C-17), 142.97 (C-1),134.37 (C-15, C-16), 133.81 (C-5), 132.30 (C-4, C-6), 132.22 (C-3), 131.80 (C-7), 131.24 (C-2), 127.77 (C-8), 38.91 (C-13), 37.76 (C-12), 35.80 (C-10), 33.51 (C-9) ppm.

Mass spectometry (ESI, *full scan*) m/z calculated 298.3365, found 299.13827 [M+H]^+^, 321.12000 [M+Na]^+^.

**Figure S9:**
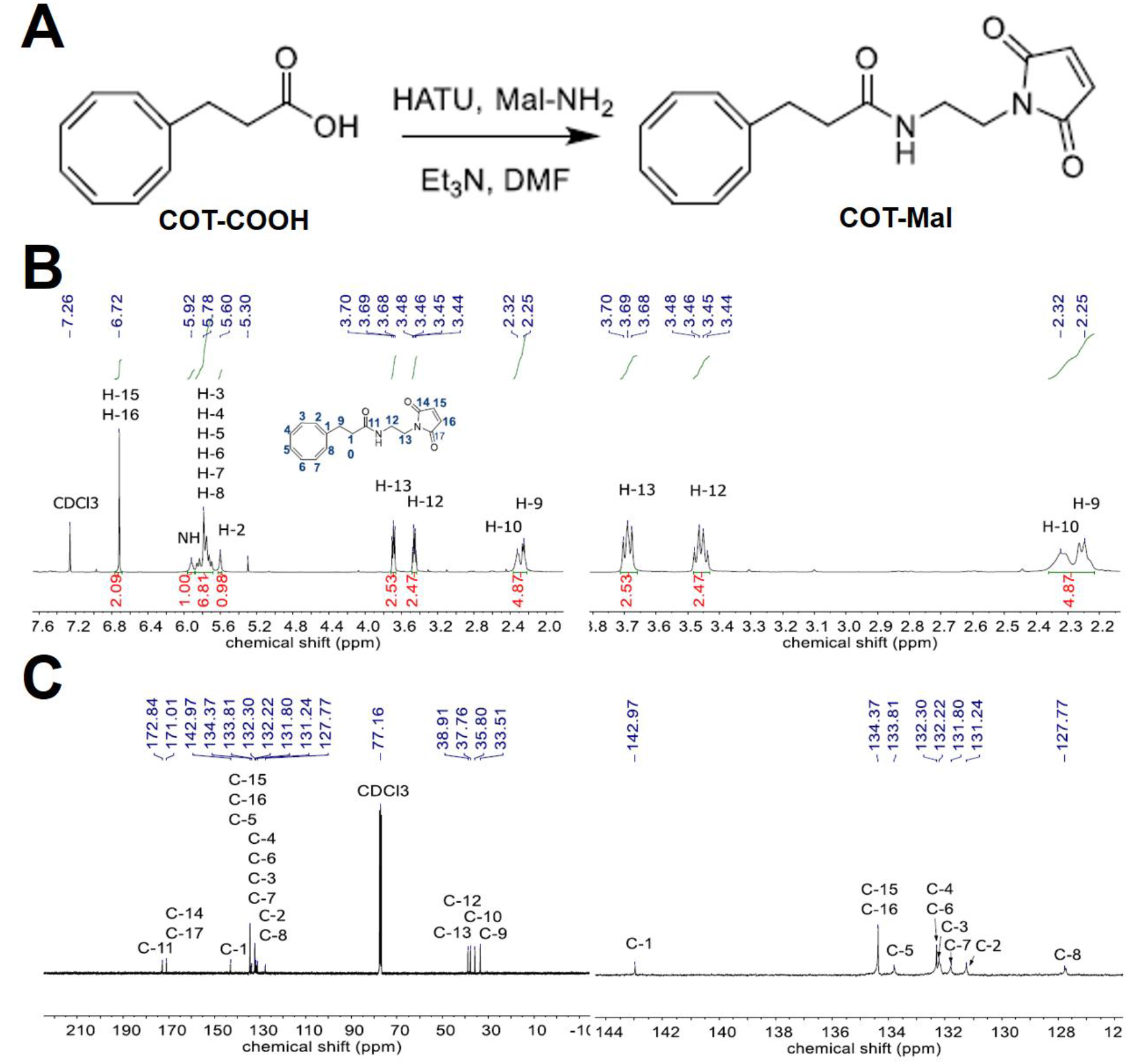
**(A)** Coupling scheme for synthesis of COT-Mal. **(B)** ^1^H spectrum and **(C)** ^13^C spectrum of COT-Mal.

